# Discovering hub genes involved in the pathophysiological impact of COVID-19 on diabetes kidney disease by differential gene expression and interactome analysis

**DOI:** 10.1101/2022.10.11.511826

**Authors:** Ulises Osuna-Martínez, Katia Aviña-Padilla, Vicente Olimón-Andalón, Carla Angulo-Rojo, Alma Guadrón-Llanos, José Carlos Rivas-Ferreira, Francisco Urrea, Loranda Calderon-Zamora

**Author notes:** Equal contribution.

## Abstract

Diabetic kidney disease (DKD) is a frequently chronic kidney pathology derived from diabetes comorbidity. This condition has irreversible damage, and its risk factor increases with SARS-CoV-2 infection. The prognostic outcome for diabetic patients with COVID-19 is dismal, even with intensive medical treatment. However, there is still scarce information on critical genes involved in the pathophysiological impact of COVID-19 on DKD. Herein, we characterize differential expression gene (DEG) profiles and determine hub genes undergoing transcriptional reprogramming in both disease conditions. Out of 995 DEGs, we identified 42 DEGs shared with COVID-19 pathways. Enrichment analysis elucidated that they are significantly induced with implications for immune and inflammatory responses. By performing a protein-protein interaction (PPI) network and applying topological methods, we determine the following five hub genes *STAT1, IRF7, ISG15, MX1*, and *OAS1*. Then, by network deconvolution, we determine their co-expressed gene modules. Moreover, we validate the conservancy of their upregulation using the Coronascape database (DB). Finally, tissue-specific regulation of the five predictive hub genes indicates that *OAS1* and *MX1* expression levels are lower in healthy kidney tissue. Altogether, our results suggest that these genes could play an essential role in developing severe outcomes of COVID-19 in DKD patients.

## 1. Introduction

Diabetes mellitus (DM) is a chronic-degenerative endocrine disease with an insidious development [1,2]. During the COVID-19 pandemic, the role of comorbidities such as DM, hypertension, cardiovascular, renal, or hepatic impairment in developing of severe COVID-19 symptoms has been highlighted [3-9]. Strong evidence suggests that the mechanisms of immunological damage in COVID-19 are like those mediated by cytokine storms, such as those presented during sepsis [10,11]. However, the knowledge regarding the mechanisms involved in developing severe disease by SARS-CoV-2 is still not explicit. Although DM has always been among the three most prevalent comorbidities among critically ill patients with COVID-19, scarce information is available about the disruptive mechanisms inducing severe illness or death.

Notably, kidney failure is a biological process closely linked to DM. Diabetes kidney disease (DKD), also known as nephropathy, is a growing health problem worldwide. This is an outcome of hyperglycemia disrupting the glomeruli, and it is defined as a progressive chronic disease, which worldwide leads the patient to dialysis and kidney transplant [12]. Regardless of insulin treatment, diabetic patients are susceptible to developing nephropathy. Around 30% −40% of patients with type 1 and 2 diabetes mellitus develop DKD. Approximately 50% of these patients will have an end-stage renal disease [13,14].

Among the processes associated with DKD are systemic and renal inflammation, where inflammatory cells such as macrophages play a crucial role, as well as biomolecules like inflammatory cytokines (IL-1, IL-6, IL-18, and TNF-alpha) and nuclear transcription factor-kappa B (NFkB). Previous reports have highlighted pathways such as Janus kinase/signal transducer and activator of transcription (JAK/STAT) as pivotal for developing this disease [15].

Currently, DKD treatment relies on using angiotensin enzyme inhibitors, angiotensin II receptor blockers, as well as statins. However, these drugs only reduce proteinuria by hemodynamic perturbations, exempting targeting the molecular factors associated with this disease’s initiation and progression. Moreover, protective biomolecules for renal diseases, including SGLT-2 inhibitors, mineralocorticoid receptor antagonists, endothelin antagonists, glucagon-like peptide-1 receptor agonists (GLP1-RA), DPP-4 inhibitors, and statins, have been tested in preclinical trials. Unfortunately, despite the significant progress achieved in clinical research, the prevalence of DKD has remained unchanged over the last 30 years [16,17]. The complex molecular events underlying DKD pathogenesis contribute to the limited treatment options.

Currently, there is no cure for diabetic glomerulosclerosis and diabetic kidney disease. Treatments are lifelong, and patients are at high risk of developing renal artery stenosis. Moreover, patients with DKD are more susceptible to developing anemy. In addition, the interstitial compartment lesion of DKD may play a vital role in the progressive fibrosis event that contributes to endogenous erythropoietin production in response to increasing levels of hypoxia recognized by this organ. This is a substantial risk enhancer for diabetes patients with a higher probability of developing left ventricular hypertrophy or heart failure [18].

Moreover, patients with diabetes are also predisposed to higher rates of kidney stone formation. Calcium-containing and uric acid nephrolithiasis kidney stones are more frequent in diabetes. Insulin resistance leads to increased levels in acidic urine. Hence, uric acid stone formation is higher in this population. This enhanced risk has also been noticed in patients with cardiometabolic syndrome and obesity. Due to gluconeogenesis and glycogenolysis impairments, patients with diabetes and kidney disease are highly susceptible to hypo and hyperglycemia [19,20]. Kidney failure is a life-threatening condition and patients suffering from this disease are a highly susceptible population.

Remarkably, since the beginning of the COVID-19 pandemic, it has been observed that those comorbidities, including the development of diabetes chronic kidney disruptions, are associated with severe symptoms in patients with SARs-CoV-2 variants [21-24]. There are recent literature describing SARS-CoV-2 infections in diabetes and DKD; for detailed review [23,25]. This medical literature points out that patients with both pathologies (DKD and COVID-19) develop multiple pathophysiological mechanisms between the lung and kidney organs.

A higher mortality rate of COVID-19 was also reported in patients with end-stage renal disease and kidney transplant recipients than in those who does not. Additionally, the Sars-CoV-2 infection relies on using angiotensin-converting enzyme 2 as an input receptor for tubular epithelial and lung epithelial cells [26,27]. Moreover, recent studies have determined that IL-6, IL-1β, and TNF-alpha cytokines participate in the cytokine release syndrome in patients with COVID-19, producing as an outcome renal inflammation and cardiovascular disorders [26,28].

In 2020, Wu et al. reported the clinical manifestations of forty-nine hospitalized hemodialysis patients infected with COVID-19 who developed pneumonia. Overall, the disease course was severe in patients with kidney failure. They conclude that patients on hemodialysis with COVID-19 were at higher risk of death [29].

Other studies suggest that among the complications that occur in hemodialysis, patients with non-invasive ventilation develops acute respiratory distress syndrome (ARDS), shock, acute cardiac injury, and arrhythmia, which were less frequent in patients without dialysis [20,23]. These observations suggest that atypical clinical presentations of COVID-19 develop in hemodialysis patients. Hence, there is a current need to implement a systematic approach to diagnose atypical cases of COVID-19. For instance, atypical symptoms, such as fatigue, diarrhea, and loss of appetite also present in other diseases complicating the diagnosis of COVI-19 [23,29,30].

Different studies have reported that dyspnea and high basal creatinine are more frequent in kidney transplant patients than in non-transplant patients [20,29,30]. Kataria *et al* reported management considerations for patients with COVID-19 who need a kidney transplant, with the aim of reducing the spread of the virus in donors, recipients, and medical staff [23,31].

The study published by *Mourad et al* shows that viral pneumonia occurred in 96% of hospitalized patients, of which 39% received mechanical ventilation and 21% received renal replacement therapy. Notably, 28% of kidney transplant patients and 64% of intubated patients died [20]. In this context, building guidelines for managing and considering atypical sing and symptoms for the differential diagnosis in kidney transplant and dialysis patients to reduce the complications during the procedure [20,32].

Even with the new development of vaccines and their distribution worldwide, it is likely that we should consider the constant interaction with new, possibly more infectious variants as the “*new normality*” [20,33,34]. It is noteworthy that the continuing global spread of SARS-CoV-2 and the latent morbidity/mortality of COVID-19 increases with comorbidities, such as DKD. Hence, it is necessary to develop strategies for the improvement of patients’ health.

In this context, bioinformatics approaches are powerful strategies, considered one of the most promising tools to predict potential therapeutic genes with clinical relevance and the biological processes underlying complex diseases [35,36]. Recent studies have identified predictive genes shared among COVID-19 and multiple comorbidities by performing bioinformatics analysis. These approaches have provided insights into molecular mechanisms and the biological processes to divert the possible final result of a severe COVID-19 condition and to provide potential therapeutic targets [37-39].

In this work, we focus on transcriptomics and network analysis to determine the shared genes between DKD and COVID-19 affected pathways, emphasizing identifying those which represent hub genes involved in kidney disease complications. Our study could contribute to a better understanding of the pathophysiological impact of COVID-19 on kidney disease complications in diabetic patients. We consider that this could serve as a basis for further research in developing improved therapy for patients suffering from these health conditions.

## 2. Materials and Methods

For this study, we designed a bioinformatics pipeline that uses transcriptomic data to identify predictive hub genes involved in DKD complications derived from COVID-19 infection as depicted in Figure 1. Used codes are available at https://github.com/kap8416/Transcriptomics-Diabetes-Kidney-Disease.

**Figure 1.**
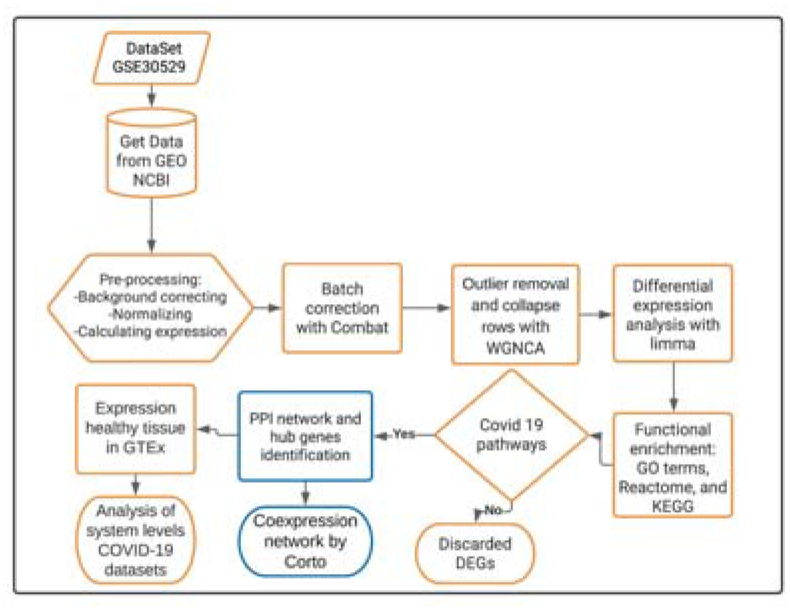
Bioinformatics pipeline for obtaining predictive hub genes involved in the pathophysiological impact of COVID-19 on DKD condition. Highlighted in yellow are the steps to perform the transcriptomics analysis, while those in blue are steps for the interactomics methodology. First, transcriptomic data was acquired from the NCBI GEO database. Datas were pre-processed, and outlier detection with WGCNA analysis was performed. Then, we obtained the DEGs and determined their potential functional roles in biological and molecular pathways. Those DEGs that overlapped among COVID-19 and DKD pathways were selected, and interactome analysis was performed using protein-protein interaction (PPI) associations. Subsequently, the potential hub genes were identified using the Maximal Clique Centrality (MCC) topological algorithm. Then, their co-expressed gene modules were determined using the “*Corto*” algorithm. Finally, hub gene expression was validated using analysis of systems levels employing Coronascape COVID-19 datasets and gene expression value comparisons in healthy tissues using the GTEx database.

### 2.1 Transcriptomic data acquisition

The microarray expression dataset (GSE30529) was obtained from the NCBI GEO database (https://www.ncbi.nlm.nih.gov/geo/). This dataset is based on the GPL571 platform and comprises human samples from patients with DKD (n = 10) and healthy (non-diabetic) controls (n = 12).

### 2.2 Differential gene expression analysis

Raw data were processed with the “*affy*” package available in Bioconductor using the function robust multi-array average (RMA) to convert an “*AffyBatch*” object into an “*ExpressionSet*” object (https://www.bioconductor.org/). Then, data normalization was performed. After that, we removed the “batch effect” using “*ComBat*” from the *sva*, and Weighted Gene Co-Expression Network Analysis (WGCNA) to identify outliers. Also, the *collapseRows* function available in the WGCNA package for collapsing gene expression of several probes in a single measurement per gene, was used. Finally, differential gene expression analysis was performed using the limma package. Log2FC > 1, and FDR < 0.05 values were considered to determine the differentially expressed genes (DEGs). Genes having significant *p-values* with positive Log2FC represent an increased expression (UP). Those with negative Log2FC values are considered downregulated (DN), while those with *p-values* above 0.05 do not have changes between stages (NC).

### 2.3 Functional enrichment analysis for DEGs in the DKD dataset

The functional enrichment was assessed using the Gene Set Enrichment Analysis (GSEA) of DEGs. Enrichment includes all the Gene Ontology (GO) terms and the Kyoto Encyclopedia of Genes and Genomes (KEGG), and Reactome pathways. The cluster profiler R package [40] was employed for data analysis and SRplot server (http://www.bioinformatics.com.cn/en?keywords=pie) for data visualization. Further, we considered focusing only on 42 genes shared in both DKD and COVID-19 pathogenetic processes.

### 2.4 Functional enrichment analysis for the shared pathogenic DEGs

Functional enrichment analysis was performed for the DEGs using KEGG and Reactome pathways, including the GO terms for biological processes. Then, the ClueGO (version 2.5.7) module of the Cytoscape software (version 3.9.0) was used to examine the inter-relational pathway analysis of the significantly enriched functions to identify the most significant genes [41]. We identified up and downregulated pathways using the top 200 DEGs. Then, the 42 overlapped genes were considered to perform the same analysis with a kappa score of 0.4, *pvalue* < 0.05. For statistical tests, hypergeometric two-sided and Benjamini–Hochberg methods were employed [41]. Finally, the Cluster profiler R package was employed for data analysis and visualization [40].

### 2.4 PPI network and identification of hub genes

The list of 42 DE-human genes previously identified in the shared pathogenetic process were analyzed to deep insight into their interatomic roles. Network analysis and visualization were performed using the Cytoscape version 3.9.0. STRINGDB platform (https://string-db.org/) was used to obtain physical and functional experimental validated data. For this study, high confidence scores (0.7) and no additional interactors were filtered, keeping only experimental, co-expression and database characterized interactions. Each node in the PPI network represents a protein whereas edges are interactions and connections between them. Finally, the predictive hub genes were obtained using the CytoHubba module selecting the top five nodes employing the topological algorithm of Maximal Clique Centrality (MCC) [42].

### 2.5 Deconvolution of the Gene Regulatory Network (GRN)

In order to identify co-expression modules of the five hub genes, the “*Corto*” algorithm with default parameters was used for the GRN inference. This package is freely available on the CRAN repository. The “*Corto*” algorithm is implemented as a co-expression-based tool that infers GRNs using as an input a TFs list with their targets and an expression matrix data set [43]. In brief, Corto uses a combination of *Spearman* correlation and Data Processing Inequality (DPI), adding bootstrapping to evaluate the significant edges by removing indirect interactions. We used the DKD expression matrix and a human-TF list as input. For this analysis, a *p-value*= 1 × 10-8 was used as a cut-off, and 100 bootstraps were carried out. Subsequently, an output file containing an inferred enriched GRN was obtained. For network visualization, the Cytoscape environment was used to identify the co-expressed gene modules of each hub gene [44].

### 2.6 Functional enrichment analysis for co-expressed DEGs in the GRN

The functional enrichment analysis was assessed using the GSEA of each of the five previously identified modules of the hub genes co-expressed DEGs. Functional enrichment considered all the GO terms and the KEGG and Reactome pathways. The cluster profiler R package was employed for data analysis [40]. For data visualization the parent terms were depicted in the GRN using the Biorender environment (https://biorender.com/).

### 2.7 *In silico* validation *of expression of the predictive hub genes in Coronascape DB*

Gene expression data was accessed at the Coronascape repository [45]. Coronascape is a resource for the analysis of systems-level datasets. The top statistically overlapped COVID reference lists datasets were analyzed against our gene list (5 hub-DEGs). DEGs conserved among the COVID databases were identified and visualized in Circle Plot using the Circos visualization tool (https://genome.cshlp.org/content/early/2009/06/15/gr.092759.109.abstract) for comparative genomics analysis.

### 2.8 In silico validation of expression of the predictive hub genes in GTEx database

Gene expression data was accessed at the GTEx repository (https://www.gtexportal.org/). GTEx is a resource for the analysis of tissue-specific level datasets. The five hub genes were tested for multi-gene query expression in kidney, lung, heart, pancreas, and whole blood healthy tissues. Data was visualized in a heatmap plot. Then, single-cell data from lung tissue was accessed to determine the specific cell expression of the hub genes. Data was visualized in pie charts.

## 3. Results

### 3.1. Identification of Differentially Expressed Genes across Diabetic Kidney Disease

First, we performed raw data normalization using the No GSE305029 Affymetrix microarray dataset. This step allowed reducing false positives, as depicted in Figure 2A. Later, principal component analysis (PCA) showed clustered distances among each renal tubule sample, including those belonging to the disease condition (DKD) and those to the control group (Figure 2B). After this step, one *batch* was observed for the sample GSM757034 belonging to the control group. This sample was removed for further dataset proccesing. After that, we performed WGCNA. This approach is a widely used data mining and exploratory method. It allowed us to identify clusters and outliers to discard samples that did not cluster with their corresponding group (disease or control). Following this step, we removed sample GSM757027. After that, two separate clusters according to their patient phenotype are depicted in Figure 2C and employed further for the transcriptomics analysis. By performing gene expression analysis, we identified a total of 13,309 genes in the microarray samples. Nine hundred ninety-five out of 13,309 genes are DEG in the disease samples when compared to kidney tubules. Remarkably our results showed that DEGs are significantly upregulated. Around ∼63% of them (632 genes) are induced, while only 363 were undergoing down-regulation of their expression (Figure 2D, Table S1).

**Figure 2.**
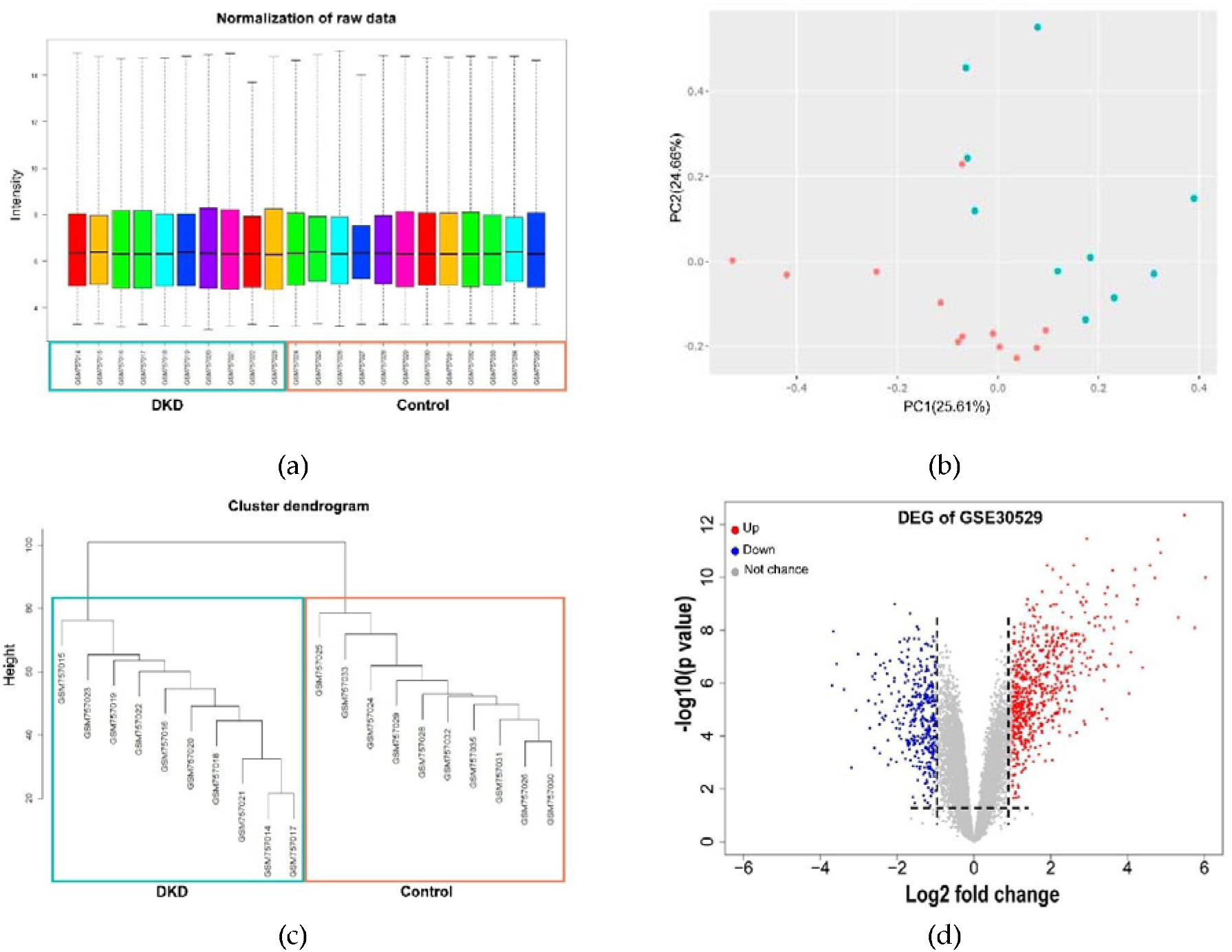
Pre-processing of GSE30529 dataset raw microarray data and differential expression analysis. (a) Boxplot of normalized raw data contrasting human patient healthy and DKD samples from kidney tubule. (b) PCA plots depict similarities and differences among DKD and control samples after data normalization. (c) WGCNA clustering represents 20 gene modules and two clusters. (d) Volcano plot of DEG, red (up) and blue (down) colored dots indicate the DEGs, while the grey dots represent genes without expression changes (NC) among DKD and control samples.

The top twenty deregulated genes are depicted in **Figure 3**. The top ten upregulated genes identified are *IGHA2* (Log2FC= 6.027, fdr=9.88E-11), which encodes immunoglobulin receptor binding. This gene is involved in glomerular filtration, and its related pathway is the inflammatory response pathway. Followed by *IGHGP* (Log2FC= 5.749, fdr=8.16E-09) a pseudogene accordingly non-functional gene. Then, *LTF* (Log2FC= 5.482, fdr= 4.32E-13) is a member of the transferrin family of genes with antimicrobial, antifungal, antiparasitic, and antiviral activity against both DNA and RNA viruses, including activity against SARS-CoV-2. Meanwhile, *IGLC2* (Log2FC= 5.320, fdr=3.27E-09) is a gene involved in several processes, including activation of the immune and defense responses to other organisms and phagocytosis. Also, it is related to the network map of the SARS-CoV-2 signaling pathway. Followed by *JCHAIN* (Log2FC= 4.861, fdr=1.17E-11). This gene participates in IgA binding activity and protein homodimerization activity. It also contributes to several processes, including glomerular filtration and positive regulation of respiratory burst. Also, we identified *CXCL6* (Log2FC= 4.793, fdr= 3.73E-12), a member of the *CXC* chemokine family with an antibacterial action against gram-positive and gram-negative bacteria, *LYZ* (Log2FC= 4.710, fdr= 1.02E-10) a gene encoding a protein with antimicrobial activity in human milk and other tissues such as lungs and kidneys. *C*3 (Log2FC= 4.587, fdr= 3.53E-11) is a protein-coding gene that plays a central role in the activation of the complement system. This gene encoded a peptide that modulates inflammation and possesses antimicrobial activity. *IGHM* (Log2FC= 4.390, fdr= 2.60E-07) is a gene that encodes the C region of the heavy mu chain, which defines the IgM isotype. *ALOX5* (Log2FC= 4.261, fdr= 6.81E-10) is a gene member of the lipoxygenase gene family and plays a dual role in the synthesis of leukotrienes from arachidonic acid. Notably, leukotrienes are important mediators of several inflammatory and allergic conditions. (**Figure 3, Table S1**).

**Figure 3.**
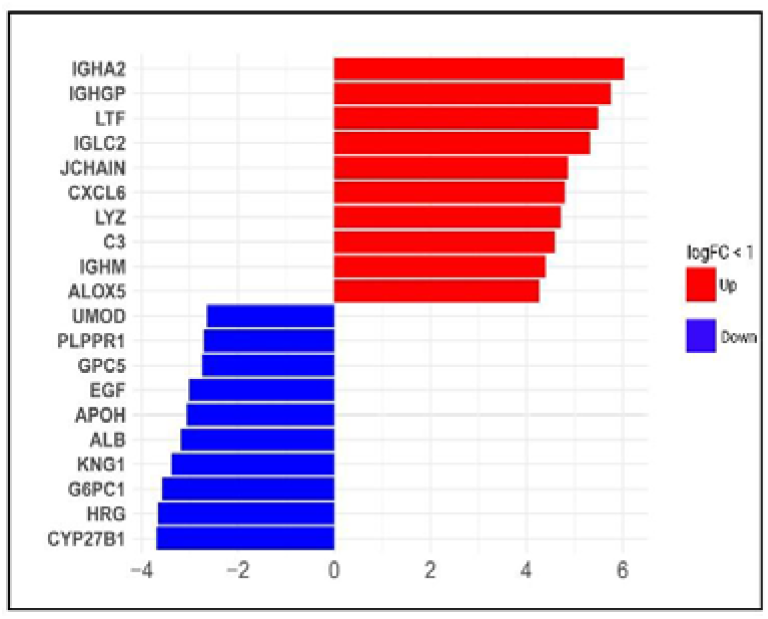
Top upregulated and downregulated DEGs in diabetic kidney disease dataset. Bars show the top upregulated (red) and downregulated (blue) genes, according to Log2FC values.

Opposite, the top ten downregulated DEGs listed are the following. *CYP27B1* (Log2FC= −3.693, fdr= 1.24E-06) encodes a protein member of the cytochrome P450 superfamily of enzymes and is related to the steroid metabolism pathways. Followed by *HRG* (Log2FC= −3.661, fdr= 1.09E-08) a gene encoding a histidine-rich glycoprotein located in plasma and platelets. The physiological function has not been determined. However, the encoded peptide displays antimicrobial activity against *C. albicans, E. coli, S. aureus, P. aeruginosa, and E. faecalis* microorganisms [46]. Followed by *G6PC1* (Log2FC= −3.575, fdr= 1.87E-07) a gene that catalytic-subunit-encoding glucose-6-phosphatase enzyme involved in glucose homeostasis, *KNG1* (Log2FC= −3.383, fdr= 1.76E-06) that uses alternative splicing to generate two different proteins-high molecular weight kininogen (*HMWK*) and low molecular weight kininogen (*LMWK*). *HMWK* is essential for blood coagulation and the release bradykinin [47,48]. Interestingly, during SARS-CoV-2 infection an increase in bradykinin levels is associated with lung injury and inflammation [49]. Meanwhile, *ALB* (Log2FC= −3.188, fdr= 0.001) a gene that encodes the most abundant protein in human blood. This protein acts to regulate blood plasma colloid osmotic pressure and as a carrier protein for a wide range of endogenous biomolecules. Additionally, this protein encoded a peptide that is an endogenous inhibitor of the *CXCR4* chemokine receptor. *APOH* (Log2FC= −3.067, fdr= 1.09E-05) a gene coding by apolipoprotein H, a component of circulating plasma lipoproteins. *EGF* (Log2FC= − 3.017, fdr= 8.05E-08) this gene encodes a member of the epidermal growth factor superfamily. This protein acts as a potent mitogenic factor that plays an important role in numerous cell types of growth, proliferation, and differentiation. *GPC5* (Log2FC= −2.746, fdr= 4.63E-06) a protein-coding gene related with SARS-CoV-2 Infection pathway is also identified as downregulated. Besides, *PLPPR1* (Log2FC= −2.711, fdr= 5.00E-05) a gene of member plasticity-related gene (*PRG*) family (Log2FC= −1.61E-08, fdr= 5.64E-07) which is the most abundant protein in mammalian urine under physiological conditions, and its urine excretion may defend against urinary tract infections caused by uropathogenic bacteria (www.genecards.org).

Altogether, our results from the differentially expression analysis of DKD samples show that the most upregulated genes are a group of immunoglobulin receptors that could be participating in the activation of the immune and defense responses to other organisms and phagocytosis. Meanwhile, the downregulated genes are a group of biomolecules associated with mitochondria and other cell compartments metabolism, particularly lipoproteins.

### 3.2. DEGs in DKD are significantly enriched in cell and immune responses and COVID-19 affected pathways

To delve into the roles of the DEGs, we performed a functional enrichment analysis, Figure 4. Our results indicates that the DEGs are highly enriched in the following immunological biological processes: leukocyte cell-cell adhesion (count = 65, p = 5.10E-28), leukocyte proliferation (count = 62, p = 4.78E-29), adaptive immune response based on somatic recombination of immune receptors built from immunoglobulin superfamily domains (count = 61, p = 3.48E-25), positive regulation of cell activation (count = 60, p = 1.52E-22), regulation of leukocyte cell-cell adhesion (count = 59, p = 4.09E-25), extracellular matrix organization (count = 59, p = 1.97E-24), extracellular structure organization (count = 59, p = 2.22E-24), positive regulation of leukocyte activation (count = 59, p = 7.37E-23), regulation of T cell activation (count = 57, p = 1.78E-23), humoral immune response (count = 57, p = 1.95E-22), regulation of leukocyte proliferation (count = 55, p = 1.10E-27), lymphocyte proliferation (count = 55, p = 5.00E-25), mononuclear cell proliferation (count = 53, p = 7.48E-25), and positive regulation of lymphocyte activation (count = 53, p = 9.44E-21) (**Figure 4A, Table S2**).

**Figure 4.**
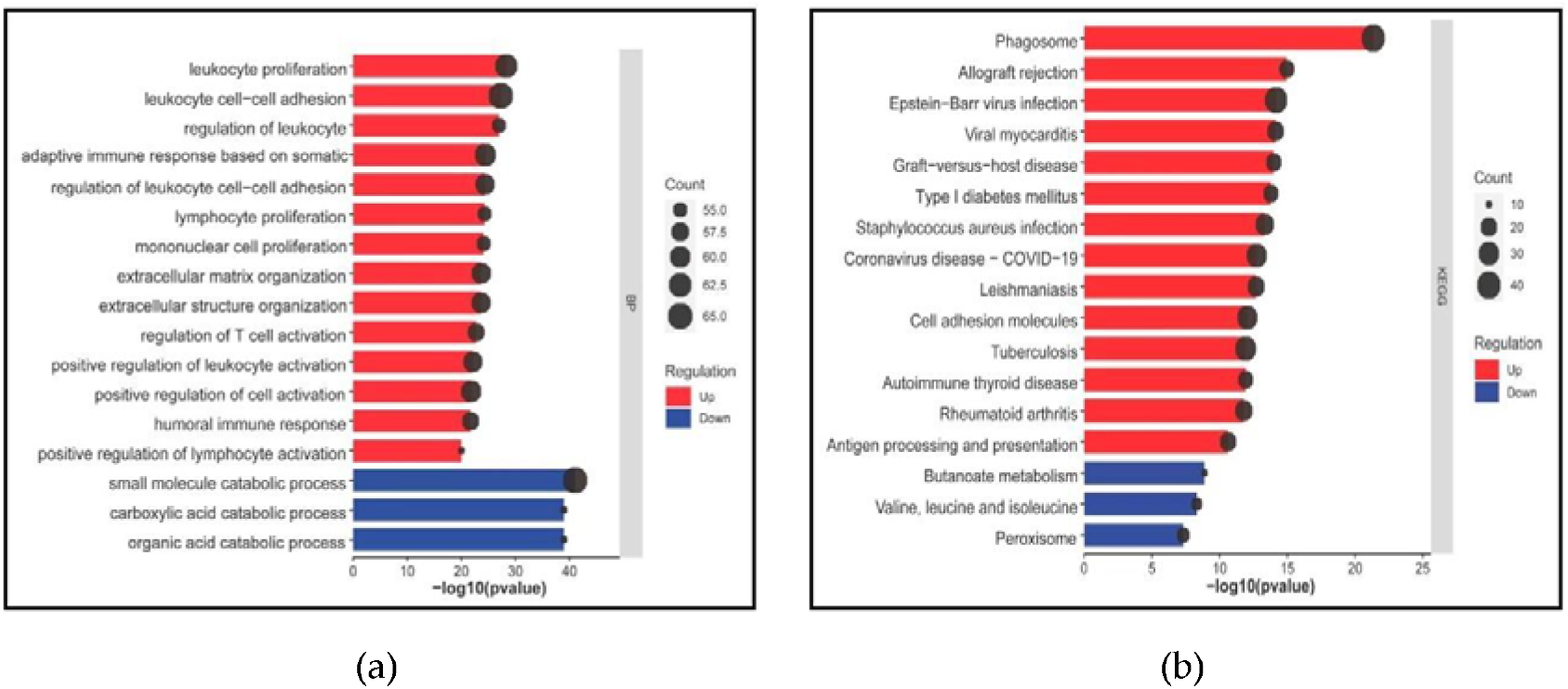
Functional enrichment analysis of DEGs. (a) Gene Ontology biological processes, and (b) KEGG pathways. Bars for upregulated are colored in red while downregulated are depicted in blue.

Among those, phagosome (count = 40, p = 4.57E-22), allograft rejection (count = 18, p = 1.10E-15), epstein-Barr virus infection (count = 37, p = 6.80E-15), viral myocarditis (count = 21, p = 7.51E-15), graft-versus-host disease (count = 18, p = 1.00E-14), type I diabetes mellitus (count = 18, p = 1.66E-14), *Staphylococcus aureus* infection (count = 25, p = 4.67E-14), Coronavirus disease-COVID-19 (count = 30, p = 1.82E-13), Leishmaniasis (count = 22, p = 2.03E-13), cell adhesion biomolecules (count = 30, p = 9.00E-13), tuberculosis (count = 32, p = 1.20E-12), autoimmune thyroid disease (count = 18, p = 1.21E-12), rheumatoid arthritis (count = 23, p = 1.63E-12), and antigen processing and presentation (count = 20, p =2.36E-11) are induced pathways (**Table S3**).

In contrast, small molecule catabolic process (count = 65, p = 9.17E-42), organic acid catabolic process (count = 53, p = 1.07E-39), and carboxylic acid catabolic process (count = 53, p = 1.07E-39) are repressed biological processes as well as (Figure 3B, Table S2) route blades as are butanoate metabolism (count = 10, p = 1.32E-09), valine, leucine and isoleucine degradation (count = 12, p = 4.83E-09), and peroxisome related pathways(count = 14, p = 4.81E-08) (**Figure 4B, Table S3**).

Interestingly, upregulated DEGs were implicated in autoimmune disorders and bacterial and viral infections such as COVID-19. Whereas downregulated DEGs were involved with amino acid metabolism pathways. Altogether, the most enriched pathways demonstrate the significant role of the DEGs in diabetic nephropathy, activating immunological and defense responses commonly affected in the SARS-CoV-2 scenario. Opposite, the top enriched biological processes are linked to amino acid and metabolism pathways, suggesting their role in the consequence is a profound disturbance in glycolysis and lipid and amino acid metabolism.

### 3.3. Transcriptional reprogramming of gene upregulation is a common mechanism for DKD and COVID-19 conditions

Notably, our results showed that ∼67.6% of DEGs in DKD samples are upregulated. From the 632 upregulated genes, 41 out of the 560 genes involved in COVID-19 pathways overlapped (**Figure 5A, Table S4**). Interestingly, the overlapped DEGs among DKD and COVID-19 affected pathways also include immune, diabetes, and other comorbidities according to their role in KEGG, Reactome, and GO terms, (**Figure 5B**).

**Figure 5.**
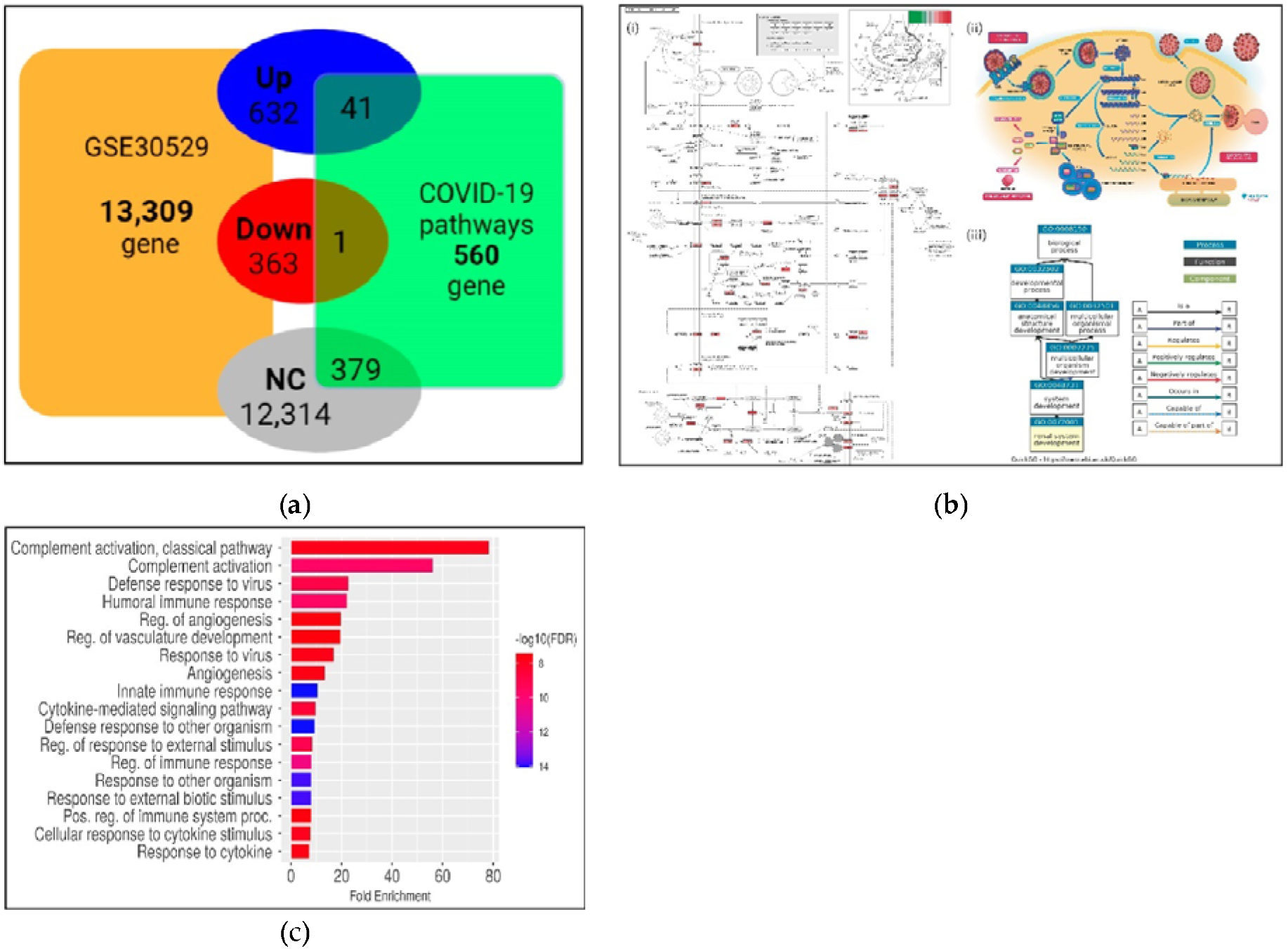
Identification of upregulated genes in DKD associated with COVID-19 pathways. (a) Diagram of overlapping DEGs among DKD and the COVID-19 pathways. The intersection of forty-one genes in the COVID-19 pathway is shown. (b) (i) The KEGG pathway map hsa05171 is involved in Coronavirus disease - COVID-19. The rectangle in red color (light to dark), according to log Log2FC, indicates the upregulated DEGs overlapping DKD and COVID-19. ii) the Reactome pathway R-hsa-9694516, and iii) the GO term corresponding to renal system development. (c) Functional enrichment analysis of the shared genes among DKD and COVID-19 pathways. FDR is calculated based on a nominal p-value from the hypergeometric test. Fold Enrichment is defined as the percentage of DEGs belonging to a pathway divided by the corresponding percentage in the background. FDR reports how likely the Enrichment is by chance. Higher values are colored on a scale of red to blue. In the x-axis, Fold Enrichment indicates how drastically genes of a specific pathway are overrepresented.

Among the top fifteen genes involved in the identified pathways are the described in **Table 1**. *C3* is highly significant, this gene plays a central role in the activation of the complement system and encodes the C3a peptide, which modulates inflammation and possesses antimicrobial activity [50]. While *CFB* gene is a component of the alternative pathway of complement activation, which circulates in the blood as a single-chain polypeptide. *CASP1* plays a central role in cell apoptosis. This gene activates the inactive precursor of interleukin-1, a cytokine involved in the processes such as inflammation. *TLR7* and *TLR2* are essential in recognizing pathogen-associated molecular patterns (PAMPs) by infectious agents that mediate the production of cytokines necessary for developing effective immunity. *TLR7* participates in the recognition of single-stranded RNA viruses, which this gene is associated with COVID-19 [51]. *CXCL10, CXCL8*, and *CCL2* are cytokine genes involved in inflammatory processes. *CXCL10* is a gene key regulator of the ‘*cytokine storm*’ immune response to SARS-CoV-2 infection, and *CCL2* is associated with severe acute respiratory syndrome coronavirus 2 [52-54].

**Table 1.** Top fifteen DEGs overlapped among DKD and COVID-19 pathways. Presence: (^*^) or not (NA)

| Gene description | Gene symbol | KEGG | Reactome |
| --- | --- | --- | --- |
| Complement C3 | C3 | * | NA |
| Caspase 1 | CASP1 | * | NA |
| C-C motif chemokine ligand 2 | CCL2 | * | NA |
| Complement factor B | CFB | * | * |
| C-X-C motif chemokine ligand 10 | CXCL10 | * | NA |
| C-X-C motif chemokine ligand 8 | CXCL8 | * | NA |
| Interferon alpha and beta receptor subunit 2 | IFNAR2 | * | * |
| Interferon regulatory factor 7 | IRF7 | * | * |
| ISG15 ubiquitin like modifier | ISG15 | * | * |
| Janus kinase 1 | JAK1 | * | * |
| MX dynamin like GTPase 1 | MX1 | * | NA |
| 2'-5'-oligoadenylate synthetase 1 | OAS1 | * | * |
| Signal transducer and activator of transcription 1 | STAT1 | * | * |
| Toll like receptor 2 | TLR2 | * | NA |
| Toll like receptor 7 | TLR7 | * | NA |

Moreover, *CXCL8* plays a role in the pathogenesis of the lower respiratory tract infection bronchiolitis, a common respiratory tract disease caused by the respiratory syncytial virus (*RSV*). In comparison, *JAK1* is a member of a class of protein-tyrosine kinase. This gene is a component of the *IL6/JAK1/STAT3* immune and inflammation response and is a therapeutic target for alleviating cytokine storms. *MX1* encodes a guanosine triphosphate, a metabolizing protein that participates in the cellular antiviral response. This protein is induced by type I interferon like *IFNAR2* and antagonizes the replication process of several different RNA and DNA viruses. Another identified gene, *IRF7* encodes interferon regulatory factor 7, a member of the interferon regulatory transcription factor (IRF) family. This gene plays a role in the transcriptional activation of virus-inducible cellular genes and participates in the innate immune response against DNA and RNA viruses. While *STAT1* is a gene that can be activated by different ligands, including interferon-alpha, interferon-gamma, *EGF*, platelet-derived growth factor, and *IL6*. This protein mediates the expression of various genes and plays an essential role in immune responses to viral and other pathogens. Consequently, *ISG15* is a ubiquitin-like protein conjugated to intracellular target proteins upon activation by interferon-alpha, and interferon-beta-like *STAT1* activates interferon-alpha. *OAS1* plays a crucial role in the innate cellular antiviral response, including SARS-CoV-2 (www.genecards.org).

Then, functional enrichment analysis was performed to determine the predictive roles of the forty-one common upregulated genes. As could be expected, in accordance with the enrichment analysis of the total upregulated DEGs, the most enriched pathways are those associated with defense and immune responses, **Figure 5C**. For instance, complement activation, defense response, and humoral immune response were the top enriched terms. Moreover, other affected processes identified in the upregulated genes are clustered in angiogenesis and vasculature development events. Besides, these induced genes are participating in cellular response to viruses and to cytokine stimulus.

Further, to determine their involvement in multiple molecular-biological processes, an inter-relational analysis was performed using the ClueGO module of the Cytoscape environment to obtain functionally organized pathway-terms networks. ClueGO employs kappa statistics to link the functional terms in the network. As shown in **Figure 6A**, the upregulated DEGs were mainly involved in the innate immune system, leukocyte differentiation, angiogenesis regulation, cytokine induction, extracellular matrix organization, defense response to other organisms, and regulation of peptidase activity. Meanwhile, the inter-relational analysis of the downregulated genes reveals that the most influenced connected pathways are associated with fatty acid and lipid metabolic processes. However, other essential routes are highlighted in this analysis even though they are not highly associated with each other—for instance, anion transport, response to insulin, and renal system development, **Figure 6C**.

**Figure 6.**
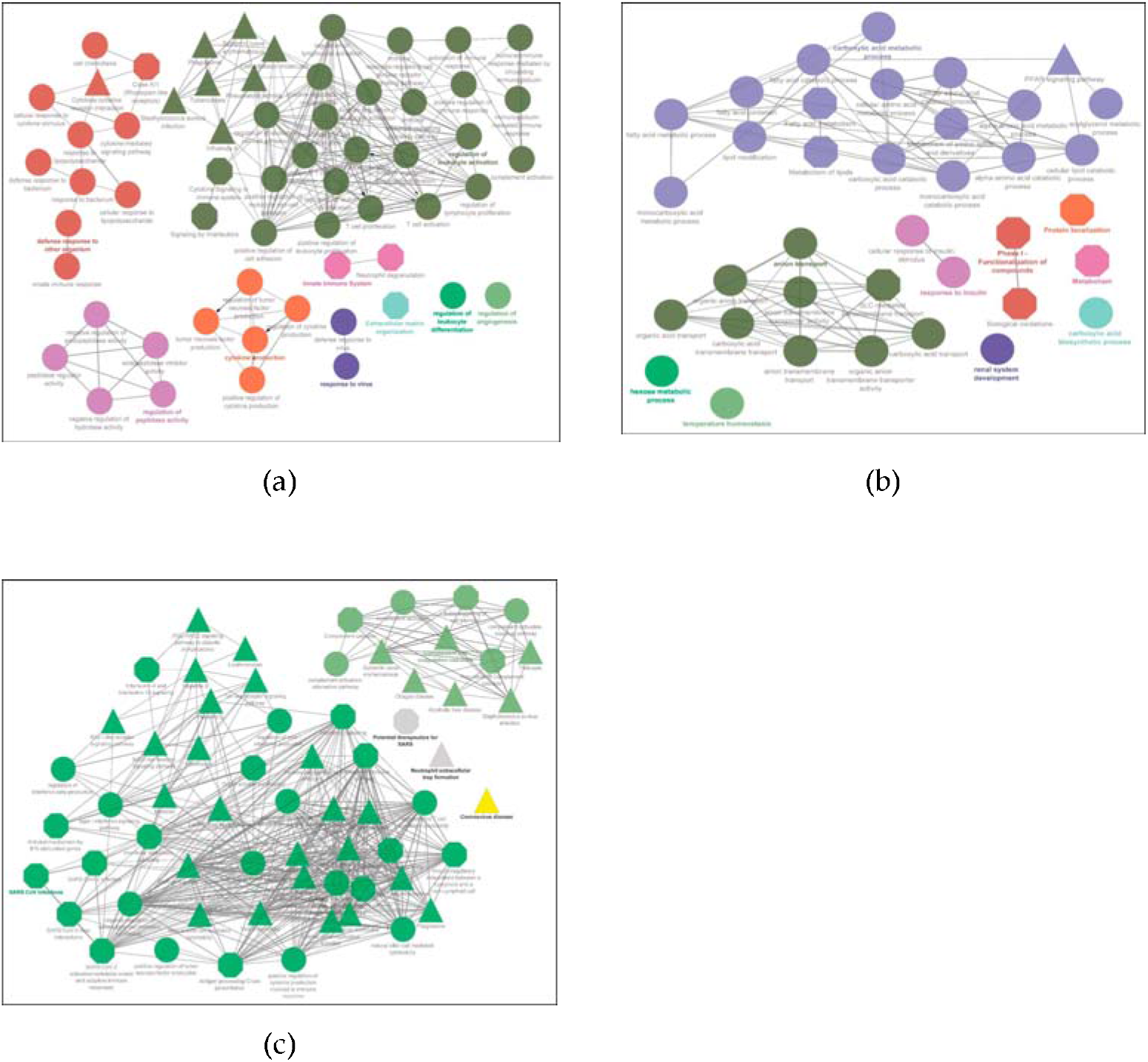
Network analysis of the functional role of deregulated genes in diabetic kidney disease. (a) Inter-relational pathway enrichment analysis is shown for the upregulated DEGs in DKD samples; (b) for the downregulated DEGs in DKD; (c) for the 41 DEGs overlapping DKD and COVID-19 pathways. Circle represents GO terms, triangle and octagon represent KEGG and REACTOME, respectively. While green, orange, purple, and pink color represent pathways.

To delve insight into the genes participating in the inter-relational pathways, we identify the potential biological involvement of the upregulated DEGs in KEGG, Reactome, and GO metabolic pathways, **Figure 7**. Notably, our results show that the five enriched terms in KEGG are coronavirus disease - COVID-19 (count = 30, p = 4.95E-38), followed by complement and coagulation cascades (count = 10, p = 1.10E-11), type I diabetes mellitus (count = 5, p = 2.35E-06), lipid and atherosclerosis (count = 8, p = 1.11E-05), and AGE-RAGE signaling pathway in diabetic complications (count = 5, p = 1.48E-04).

**Figure 7.**
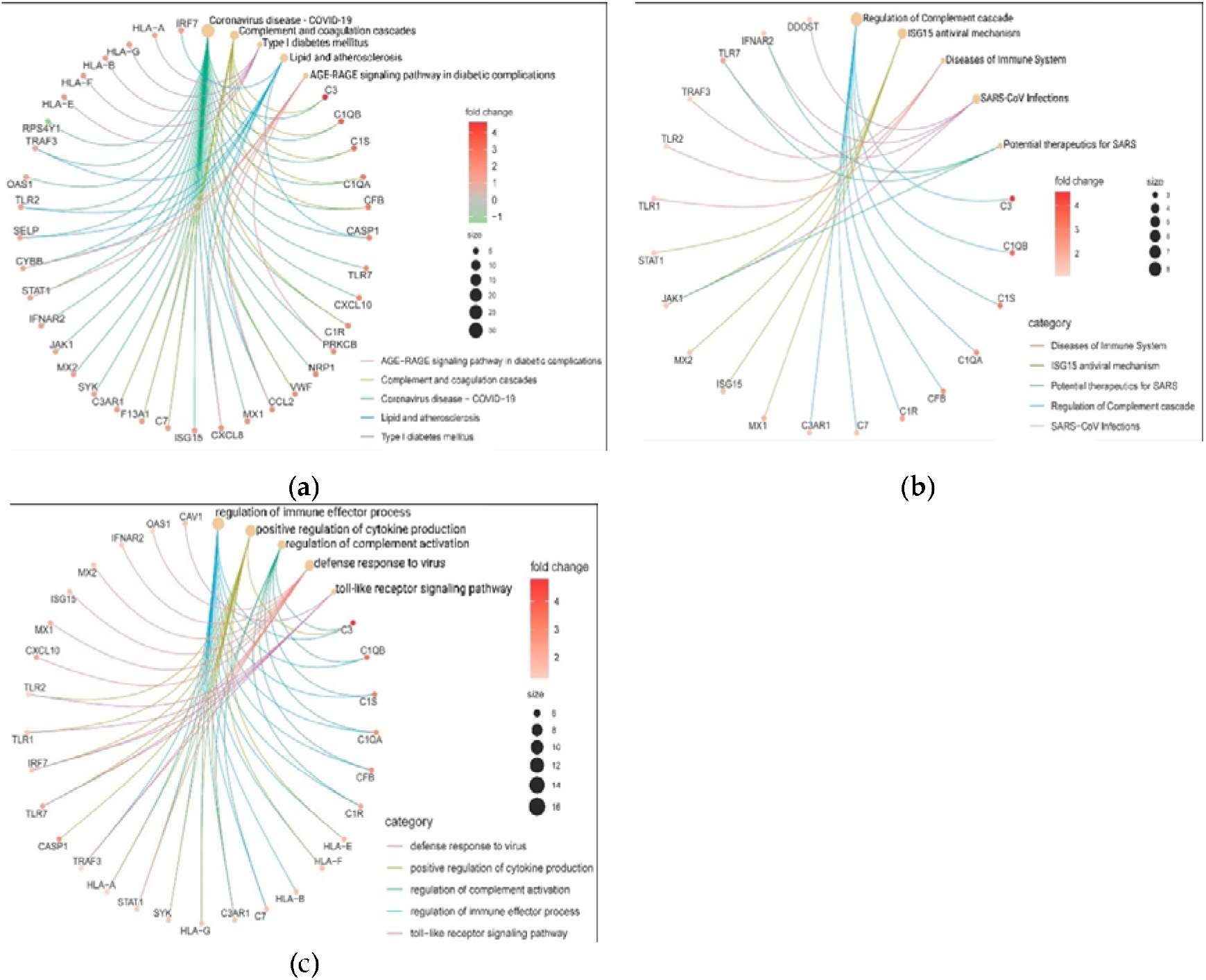
Category cnetplot depicts the linkages of upregulated genes and the biological pathways as a network. Red (light to dark) nodes indicate the Log2FC values, while yellow nodes indicate the functional enriched pathways whose size represents the number of genes that are DE in each of them.

We identified that *C3, C1QA, C1QB, C1S, CFB, CXCL10*, and *VWF* were associated with coronavirus disease and complement and coagulation cascades. While *CYBB, STAT1*, and *PRKCB* are shared for coronavirus disease - COVID-19 and AGE-RAGE signaling pathway in diabetic complications. Also, coronavirus disease-COVID-19 and lipid and atherosclerosis have connections through *TRAF3, TLR2, SELP*, and *CASP1* deregulated molecules. Finally, *CCL2* and *CXCL8* genes are related in three pathways, Figure 7a. Meanwhile, for the Reactome pathways analysis the five enriched pathways are regulation of complement cascade (count = 8, p = 9.55E-12), *ISG15* antiviral mechanism (count = 5, p = 9.44E-06), diseases of immune system (count = 3, p = 1.15E-04), SARS-CoV infections (count = 4, p = 2.78E-03), and potential therapeutics for *SARS* (count = 3, p = 4.14E-03). Being SARS-CoV infections and potential therapeutics for SARS connected by *IFNAR2, TLR7*, and *JAK1* genes. Moreover, *JAK1* gene is also present in the ISG15 antiviral mechanisms, Figure 7b. Meanwhile, the interactions within the biological process are regulation of immune effector process (count = 16, p = 1.18E-12), positive regulation of cytokine production (count = 13, p = 1.20E-11), regulation of complement activation (count = 8, p = 1.81E-010), defense response to virus (count = 10, p = 2.48E-10), and toll-like receptor signaling pathway (count = 6, p = 1.18E-06). Hence, *C1QA, C1QB, C1S, C1R, CFB*, and *C7* are related to the regulation of the immune effector process and regulation of complement activation. *C3* and *C3AR1* are also participating in the positive regulation of cytokine production. While *HLA-A, HLA-E, HLA-F, HLA-G*, and *SKY* are involved in regulating the immune effector process and positive regulation of cytokine production. Moreover, *TLR1* and *TLR2* are involved with the positive regulation of cytokine production and toll-like receptor signaling pathway as well *IRF7, TLR7, TRAF3*, and *STAT1* are present in more biological processes, **Figure 7c**.

Our results show that ∼67.6% of DEGs in DKD samples are upregulated. Around 97% (41/42) of the DEGs associated with COVID-19 pathways are upregulated. Those biomolecules participate in crucial biological and metabolic processes, including complement and coagulation cascades, lipid and atherosclerosis, AGE-RAGE signaling pathway, and positive regulation of cytokine production. Notably, according to the Reactome database, we characterize some of those induced biomolecules as potential therapeutic targets for SARS-CoV-2 infection. In summary, our transcriptomic and functional enrichment analysis suggests that transcriptional reprogramming of gene upregulation is a common mechanism in DKD, and COVID-19 conditions linked to potential pathways involved in the development of other comorbidities.

### 3.4 Interactome analysis and hub genes identification

The PPI network analysis is a useful method for identifying the hub genes playing a critical role in complex disease interactomes. The PPI networks for up-DEGs are shown in **Figure 8a**. In the predicted PPI networks for the 42 up-DEGs, an operation with a network scoring cutoff of 0.7 was applied using the MCODE plug-in of the Cytoscape environment. It resulted in seventeen DEGs with the highest interactions in the network. Among those seventeen most relevant genes, we find the following *TLR2, TLR7, CXCL10, TRAF2, MX2, JAK1, IGNAR2*, and a cluster of HLA family proteins. Then, the five prominent upregulated hub genes, namely *STAT1, MX1, ISG15, IRF7, and OAS1*, were identified and exhibited protein-protein-validated experimental interactions with a high confidence score of 0.7.

**Figure 8.**
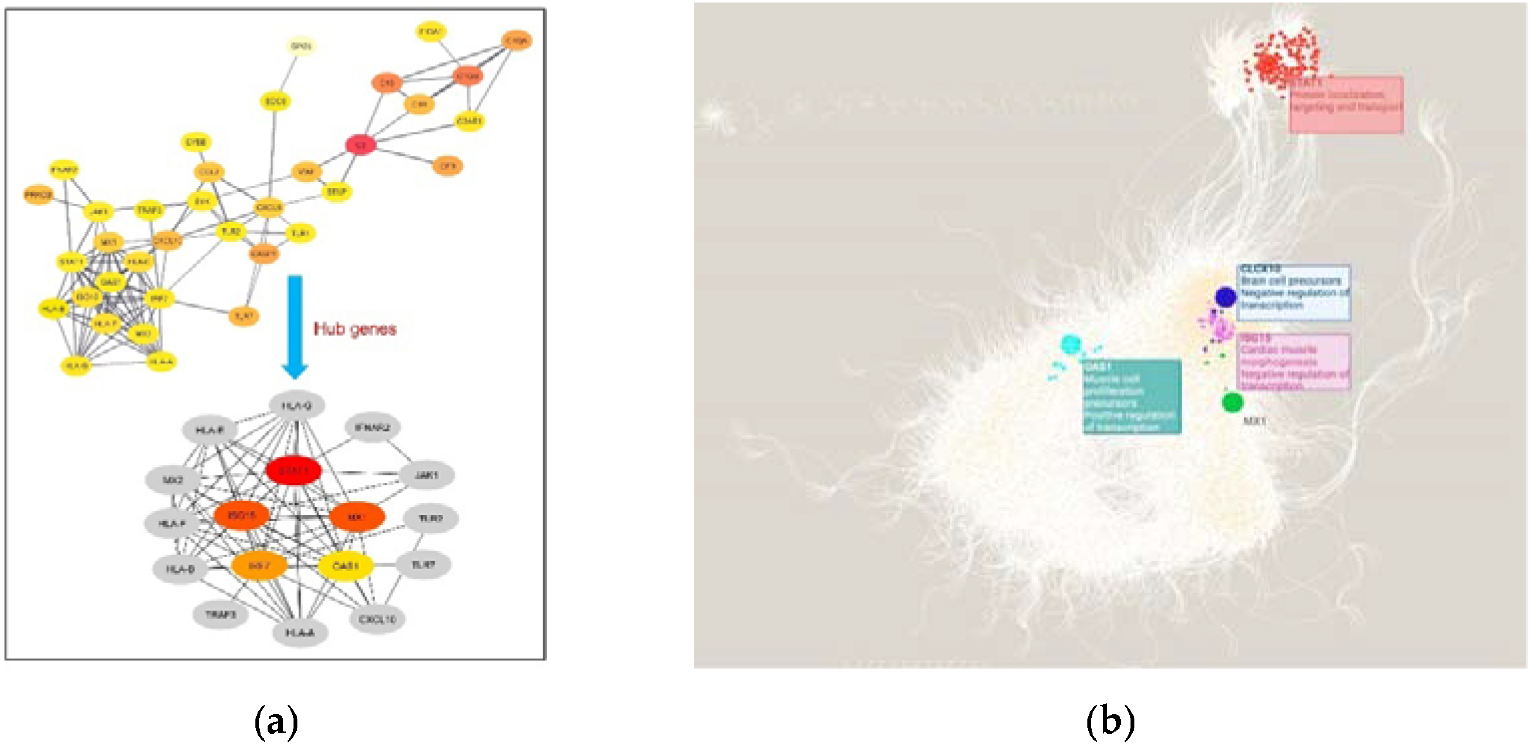
Interactome analysis of shared DEGs among DKD and COVID-10 pathways. (a) PPI network of DEGs and their closest interactors. Edges represent experimental validated interactions among proteins with a high score value (0.7). Hub genes were retrieved from cytoHubba and are shown according to score nodes (from yellow to red); (b) Interactome of the five identified hubs genes. Each node represents a gene, and the edges between nodes represent regulatory interactions among genes. The big, highlighted nodes with specific color represent each of the hub genes, while nodes sharing the same color surrounding these represent co-expressed gene modules. Enriched biological processes of each module are depicted in the same color as well. *STAT1* (red); *OAS1*(turquoise); *ISG15* (pink); IRF7 (navy blue); *MX1* (green).

When analyzing the interactome of DEGs from the GRN and their regulons, a total of 12,131 nodes, and 160, 110 interactions were obtained, **Figure 8b**. When comparing DKD samples against healthy tissue, our results showed that the *STAT1* gene module has 117 interactors, followed by the *IRF7* gene module with 43 interactions, *ISG15* presents 34 co-expressed genes, *OAS1* has 12 genes, and in the *MX1* gene module, five genes are co-expressed. Functional enrichment analysis determined that the *STAT1* gene module is associated with protein localization, targeting, and transport, while *IRF7* co-expressed genes participate in signaling by interleukins and negative regulation of immune response. *ISG15* is related to cardiac muscle, while the *OAS1* module participates in muscle cell proliferation and positive regulation of transcription, and *MX1* co-expressed genes do not present specific functional enrichment terms (Table S5).

Overall, our interactome analysis results show that the five hub genes identified are co-expressed in modules with other genes participating in the events associated with heart disease, virus immune response, and genetic regulation at the transcriptional and post-transcriptional levels.

### 3.5 In-silico validation of hub genes across COVID-19 and healthy tissue datasets

Finally, we performed in silico validation analysis to determine the expression profiles of the hub genes obtained by the interactome analysis, **Figure 9**. First, we employed the Coronascape Database, which represents a compendium of large-scale studies of transcriptomics data that allows us to perform biological systems analysis. When comparing the expression of our hub genes in DKD samples against this compendium, we identified that studies that comprise the following immune cell samples: dendritic, Calu3, CD4T, CD16/CD14 monocytes, NK and A549lowMOI cells are the ones that overlapped in the expression—being *ISG15* the most conserved differentially expressed gene (8), followed by *STAT1* (7), *OAS1*(6), and *IRF2* (2), **Figure 9a**.

Then, we delve into the expression of those genes in healthy tissue linked to the potential comorbidities’ development such as the pancreas, heart, kidney, and lung, *Figure 9b*. Additionally, we included whole blood for comparison of the baseline expression levels of those biomolecules. Our results indicate that *MX1, IRF7, ISG15*, and *STAT1* possess lung tissue-specific expression. Single-cell expression testing identified that this specific expression its higher in macrophages and B and T cells, **Figure 9c**. In contrast, *IFR7* is a gene with the highest expression values in whole blood. While *OAS1* and *MX1* are the genes with the lower expression levels in pancreas, heart, and kidney healthy samples, **Figure 9b**. Hence, the in-silico validation of our hub genes in those expression datasets pointed out that *ISG15* and *MX1* are the most conserved up-regulated genes across COVID-19 datasets of immunological cells, while *OAS1* and *MX1* are the genes with lower gene expression levels in tissue-specific healthy samples.

**Figure 9.**
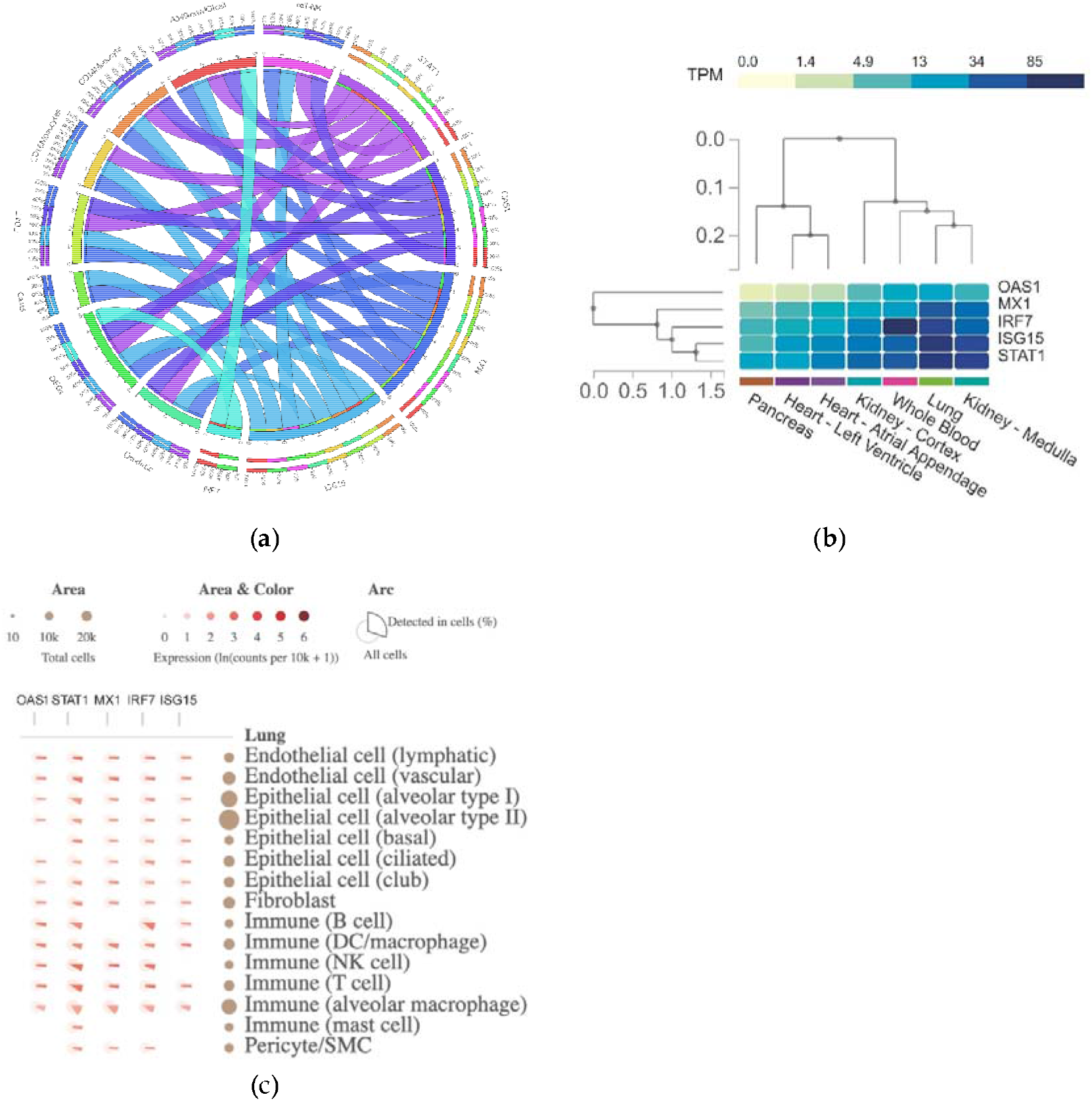
In-silico validation of gene expression of hub genes across other databases. (a) Circa plot depicts the number of shared DEGS across the 8 types of COVID-19 datasets under study. In the hemisphere, the 5 hub DEGs is represented by an arc *OAS1, STAT1, CXCL10, ISG15* and *MX1*.The inner arcs represent the distribution of the hub genes in each COVID-19 dataset, while the outside arcs represent the intersection of both hemispheres on a scale of 0–100%. Data was obtained from the Coronascape database including dendritic cells, Calu3, CD4T, CD16 monocytes, CD14 monocytes, NK cells and A549lowMOI cell samples (https://metascape.org/COVID/). (b) Heatmap comparison of hub genes healthy tissue expression and (c) lung single-cell expression using the GTEx DB (https://www.gtexportal.org).

## 4. Discussion

DM has been recognized worldwide as one of the most critical metabolic diseases because of the number of people affected and the poor quality of life they could develop. [1]. DM patients have a high rate of severe infection with the SARS-CoV-2. This fact has highlighted the impact that DM could have on the increase in fatality rates of COVID-19 in intensive care units (ICU) [21,55,56]. However, there is scarce information about the biomolecules and the biological and metabolic processes involved in worsening the clinical condition triggered by COVID-19 in diabetic patients. Hence, there is an urgent need to further our knowledge of deregulated genes as it will help to design better therapies against those complications.

Herein, we characterize at the transcriptional and post-transcriptional levels a particular group of upregulated genes and the pathogenetic events shared for DKD and COVID-19 conditions, Figure 10. Remarkably, a group of these genes is linked to biological behavior as complement and coagulation biomolecules (*C3, C1QB, CFB, CFD, C7, C3AR1* and *VWF, F13A1*). Previous studies showed that the pathophysiological mechanism of COVID-19 is associated with the complement system [57] and the coagulation system [58]. Interestingly, recent reports have described that both mechanisms have already been reported acting as enhancers. For instance, the mechanisms described in which the proteases of the coagulation cascade activate the complement pathways [59,60], and in turn, how the products of the activation of the complement pathways could activate platelets and the coagulation system. This inter-relation could be feasible to contribute to the worsening of diabetic patients with DKD infected with the SARS-CoV-2 virus. This phenomenon has been reflected in ICU rooms, giving worse results in days of hospitalization or survival of those types of patients [61-63]. Moreover, diabetic patients have also been reported to present a decrease in the fibrinolytic process [60], which further increases the risk of coagulation with COVID-19.

**Figure 10.**
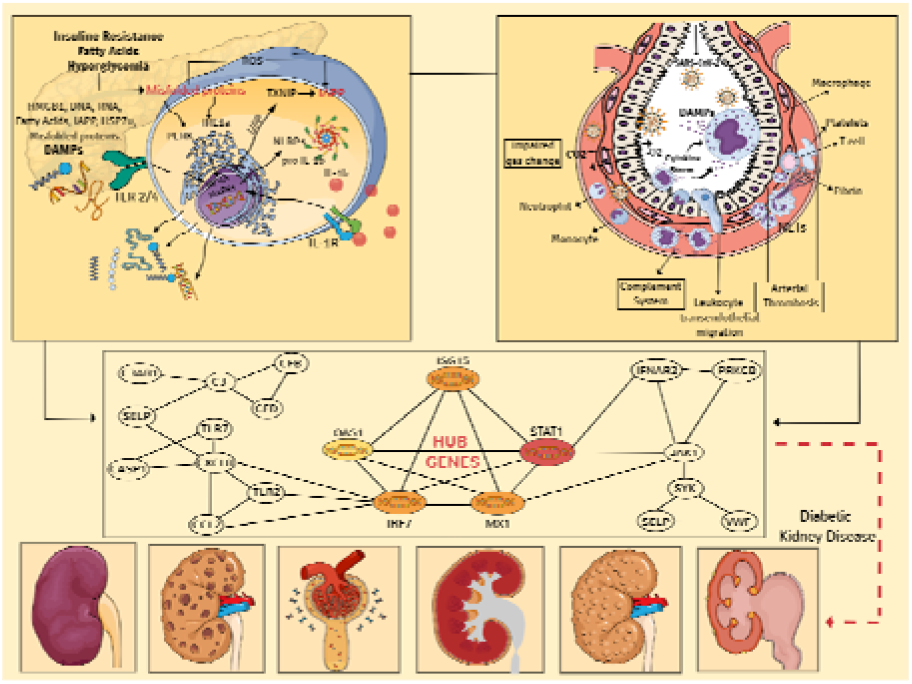
Pathophysiological impact of COVID-19 on DKD condition. (a) An illustration that simplifies the multiple processes and proinflammatory pathways that initiate in an orchestrated manner, the primary stage of the chronic inflammatory response in pancreatic cells coupled with the activation of the complement system due to the presence of DAMPs, excessive synthesis of ROS and dysfunction endothelial, contribute to the progression of DM and the development of micro and macrovascular complications in target organs, (b) pathogen-host interaction mechanisms, showing the alveolar microenvironment, where the localized inflammatory response is initiated by the presence of PAMPs and DAMPs, with the consequent endothelial activation, migrat on of immune cells, significant tissue damage, and uncontrolled release of cytokines, as well as proinflammatory interleukins (“*cytokine storm*”). (c) Predictive hub genes and their impact on the pathophysiological process of diabetic kidney disease and SARS-CoV-2 infection; (d) representation of the multiple functional and morphological alterations in renal tissue due to dysfunction of podocytes, epithelial cells, endothelial cells, and local macrophages, events that ultimately triggers the development of glomerular sclerosis, hyalinosis, deposition in the mesangial extracellular matrix and local fibrosis, alterations which together increased renal filtration, precipitation of glomerular nephrosis, acute kidney injury and progressive chronic kidney disease (CKD), among others comorbidities.

Our results also pointed out a DEG related to Neuropilin-1. Notably, like the previously described biomolecules, this gene has been associated with the worsening of COVID-19. Its role has been described to be an improvement of the entry and systemic infectivity of SARS-CoV-2 into the host. These processes would translate as pivotal factors that increase the risk of developing severe disease in diabetic patients [64]. Also, we identified the upregulation of *the IP-10 gene*. This molecule has been described as a molecular marker of severe conditions in autoimmune diseases and COVID-19. It is a genomic marker of poor prognosis or progression of the disease [65,66]. For this reason, this molecule should be taken into consideration as a possible therapeutic target if its presence is confirmed in patients with DKD.

Remarkably, other DEGs upregulated in the analyzed samples of diabetic patients are strongly related to pro-inflammatory cytokines and cell activation—for instance, *PKC, CCL2, IL-8, SYK, JAK1, IFNAR*, and *STAT1* and anti-inflammatory such as *IL-10*. As well as adhesion (*SELP*); apoptosis (*CASP1*), production of reactive oxygen intermediates (*NOX2*), and Pathogen Recognition Receptors (*PRR: TLR-2/4, TLR-7/8*). These induced genes are related to the innate immune host response against intracellular pathogens, mainly of viral type.

However, other deregulated genes are closely related to cell damage and the cytokine storm biological processes [10,67]. Particularly the PKC and SYK kinases. Both enzymes could participate in cellular biological processes such as cell activation, proliferation, and inflammation. Recently has been described their joint synergistic role in participation in monocytic cell adhesion processes [68]. In keeping with this, in the pathological context of SARS-CoV-2 that would not be beneficial due to the tissue damage that this virus incentives. As mentioned, among the upregulated genes, we also identified those related to the PRR, TLR-2/4 and TLR-7/8. These biomolecules have been described as extremely important in the defense against the virus, particularly TLR7/8 [69]. Altogether, our results suggest that the PKC/SyK actions in cells migration of the monocyte/macrophage system and the PRR participation in the overactivation of innate immune cells of the macrophage type known as macrophage activation syndrome, as well as the cytokine storm enhancer activity [70], are participating in a negative synergistic manner to increase the impact of COVID-19 in this group of the highly vulnerable patient population. This finding highlights the possibility of performing research testing the potential clinical relevance of TLRs as possible therapeutic targets. In this context, *CASP1* and *NOX2* are biomolecules closely related to PRR in innate immune defense but with a crucial role in cell damage, mainly in parenchymal or endothelial cells. *CASP1* is a molecule related to the inflammasome pathway that has a direct effect on the secretion of pro-inflammatory proteins and induces pyroptosis which is a programmed cell death by inflammation [71]. Meanwhile, the *NOX2* gene is related to the production of reactive oxygen species (ROS) [72], like *CASP1*, which if not controlled in time, end up damaging the tissue.

We also identified DEGs involved in chemotaxis and adhesion biomolecules, such as *MCP-1, IL-8*, and *SELP*. These biomolecules participate in the recruitment of immune cells, mainly effectors, like neutrophils or monocyte/macrophage type. Notwithoutstanding, their crucial role (both are very important for host protection), if they reach the parenchyma (- in the context of overactivation described for COVID-19 condition), these cells could degranulate and generate cause more damage to target organs [73,74].

Overall, our transcriptomics results suggest that the shared DEGs between DKD and SARS-CoV-2 infection represent a transcriptional reprogramming in the host response, undergoing a remarkable upregulation of key genes such as the above described, which is in accordance with previous recent studies [75,76]. For instance, the activation of antiviral biomolecule pathways such as *JAK1/STAT1* and *ISG15/MX1*, respectively. In keeping with this, *JAK1* and *STAT1* have been associated with the activation pathways of interferon secretion as for the host antiviral responses [77]. Moreover, *ISG15* and *MX1* have antiviral activity like interferon biomolecules [78-81]. It has been suggested that they may be participating in the protection against SARS-CoV-2 infection.

In our study, we also focused in analyzing the potential interactions of those deregulated molecules at their post-transcriptional level, by performing PPI and co-expression GRN analysis. We identified seventeen genes as the higher interactors among the DEGs shared in both disease conditions.Interestingly, some of the closest interactors are genes that have been previously proposed as potential therapeutics for SAR-CoV-2. For instance, *IFNAR2, IMPDH2, ITGA4, JAK1, TLR7, TUBB* have been previously identified by computational predictions and annotated in the Reactome database. All these computationally created events and entities were reviewed by Reactome curators and verified with published experimental data that indicate the existences of differences between the molecular details of the SARS-CoV-1 and SARS-CoV-2 infection pathways.

Strikingly, in our computation analysis we determined that among those genes interactors the following five are classified as hub genes *TAT1, IRF7, ISG15, MX1*, and *OAS1*. Hence, our hub genes are new molecules with promising clinical potential. Notably, those hub genes are linked to pivotal biological and metabolic pathways that are not only restricted to DKD and COVID-19 conditions, but also include heart disease, angiogenesis disruptions and other metabolic disorders. It is noteworthy that our predictive molecules could have therapeutic potential as targets. For example, in accordance to a proteome-wide genetic colocalization study performed by Anisul *et al*., *OAS1* was identified as a protein associated with the risk of COVID-19 [82]. The measured mRNA levels of *OAS1* were associated with reduced numbers in susceptibility, hospitalization, ventilation and death [83]. This study highlights that pharmacological agents that increase *OAS1* levels could be prioritized for drug development. Therefore, this evidence suggested that the induction of genes such as *OAS1* in severe COVID-19 condition could had a better prognosis in those patients [83]. Opposite, other studies showed that *OAS1* might play key roles in regulating the development of chronic kidney disease (CKD) [84]. Moreover, has been reported that the over-expression of IFN-stimulated cytokine genes such as *OAS1* and *ISG15* could contribute to systemic inflammation and as an outcome to CKD [85]. In accordance, the Gene-Disease Associations dataset describes that during acute kidney injury, the herein identified five hub genes are induced [86]. This suggests their possible involvement in leading CKD.

Another hub gene interferon-induced is *MX1*. It has been reported that this molecule is highly expressed during renal disease like lupus nephritis, and that this could be possible to cause kidney inflammation [87]. Moreover, another study supports this gene behavior. When analyzing its expression on hamster organotypic kidney cultures infected with SARS-CoV-2, it increased thirteen fold level of its baseline expression compared to the controls [88]. In this context, other genes of the IFN system that could be stimulating the signaling pathway cascades are *IRF7* and *STAT1*. Notably, *STAT1*-signaling pathway is associated with renal inflammation and renal injury [89]. Also, it has been reported an upregulation of this biomolecule in mildly to severely affected COVID-19 patients [76].

Remarkably, we observed that DKD activates several mediators in the immune response that may be participating in the excessive production of pro-inflammatory cytokines and causes some similar cytokine storm observed in COVID-19 complications [90]. Besides, conditional profiling expression in persistent cytokine in DKD may increase susceptibility to RNA viral infections like SARS-CoV-2 [91]. Its pathway begins with the binding of viral S (spike) protein to cell surface angiotensin-converting enzyme 2 (*ACE2*) and endocytosis of the bound virion.

Overall, our interactome analysis results show that the five hub genes identified are co-expressed in modules with other genes participating in molecular events associated with heart disease, virus immune response, and genetic regulation at the transcriptional and post-transcriptional levels. Our co-expression analysis highlights the regulatory role of *STAT1* in protein localization, targeting and transport. Notably, *OAS1* and *ISG15* are upregulated molecules that are both associated with the co-expression of genes linked to cardiac muscle proliferation. Our interactomics approach identified the biological possible effects of severe SARS-CoV-2 infection under DKD condition context, finding affections that are ocurring in other organs in the body. Notably, revealing linkages with cardiovascular diseases, such approaches could represent key steps towards new treatments.

In order to validate the potential of those molecules as clinic biomarkers, we performed an *in-silico* analysis taking advantage of publicly available databases. At the systems level we compared the hub genes expression against datasets in the Coronascape DB. *ISG15, STAT1*, and *MX1* are the most conserved up-regulated genes across COVID-19 datasets of immunological cells. Among immune cell samples identified to possess induction of this biomolecules in COVID-19 patients, are dendritic, CD4+T, Natural killer (NK), and CD14+CD16+ monocyte, as well as in the lung cancer cell line (calu3 and A539). Meanwhile *OAS1* is induced in all the immune cells described, to the exception of the CD16+ monocytes. In contrast, the overexpression of *IRF7* was found only for the cancer cell line A539 and in dendritic cells. NK and dendritic cells play an important role in responses to viral infections, but during Chronic kidney disease and SARS-CoV-2 infection. These immune cells decrease their circulating levels [92,93]. Opposite, the activation the intermediate (CD14™CD16+) monocytes are considered as inflammatory mediators to increase chronic conditions such as cardiovascular disease, diabetes, and DKD [94,95]. It has been reported that circulating monocytes in COVID-19 had high levels in the acute and post-acute state until up to fifteen months. Besides, CKD is associated with pro-inflammatory higher serum cytokines levels and CD4+T lymphocytes, while in hospitalized COVID-19 patients. These increased levels are associated with disease severity [96,97]. These results corroborate the important role of immune cell activation persistent in chronic kidney disease combined with the expression *ISG15, STAT1* and *MX1* could contribute to higher expression of proinflammatory cytokines potency and participate in the complication by SARS-CoV-2 infection.

Outstanding progress has been made in understanding the molecular basis of the antiviral actions of interferons. Moreover, studies have focused in delve into the strategies evolved by viruses to antagonize their actions [98-100]. Furthermore, advances on elucidating the IFN system have significantly contributed to our understanding of viral host patho-systems and disease outcomes.

Our knowledge in how molecular mechanisms underlying transcriptional and post-transcriptional events regulate the tunning in biological and metabolic pathways will contribute to translational research, from the basic research laboratory to the clinic. According to the criteria *in silico* analyzed including gene expression fold changes, hub gene scores, co-expressed modules, and their expression in healthy tissues, we proposed that the potential hub genes could have the following hierarchical order *STAT1, MX1, ISG15, OAS1*, and *IRF7* as potential prognostic or therapeutic targets. However, further studies of validation in extensive population including other potential factors, such as smoking, obesity, and geographical regions, will delineate more mechanisms underlying associations among multiple factors and COVID-19 that still need to be explored. Basic and clinical research on related genes and drugs is required to provide more strategies for preventing and treating COVID-19 in DKD condition. We hope that the new knowledge generated in this study sheds light that could lead to developing novel therapeutic intervention strategies

## 5. Conclusions

In summary, our differential expression analysis of DKD samples shows that the top upregulated genes are a group of immunoglobulin receptors that could be participating in the activation of the immune and defense responses to other organisms and phagocytosis. Meanwhile, the downregulated genes are a group of biomolecules associated with mitochondria and different cell compartment metabolism. Most DEGs in DKD samples are upregulated, being ∼ 97% of those identified to be related to COVID-19 pathways induced. Those biomolecules participate in pivotal biological and metabolic processes, including complement and coagulation cascades, lipid and atherosclerosis, AGE-RAGE signaling pathway, and positive regulation of cytokine production. Notably, those induced biomolecules include potential therapeutic targets for SARS-CoV-2 infection. Hence, transcriptional reprogramming of gene upregulation is a common mechanism in DKD, and COVID-19 conditions linked to possible pathways involved in the development of comorbidities. In addition, by performing interactomes analysis, we determine the following five hub genes *STAT1, MX1, ISG15, OAS1*, and *IRF7* and their co-expressed modules, participating in biological events associated with heart disease, virus immune response, and genetic regulation at the transcriptional and post-transcriptional levels.

Expression profile system levels analysis reveals that *ISG15* and *MX1* are the most conserved upregulated genes across COVID-19 datasets of immunological cells. Meanwhile, *OAS1* and *MX1* are the genes with lower gene expression levels in tissue-specific healthy samples. Altogether, our results pointed out that these five hub genes could play an essential role in developing severe outcomes of COVID-19 in DKD patients. The results obtained in this bioinformatics analysis could contribute to establishing future strategies using key biomolecules for clinical decision-making in the medical routine.

## Author Contributions

The following statements should be used Conceptualization, K.A.P. and L.C.Z.; methodology, K.A.P. and L.C.Z.; software, L.C.Z.; validation, K.A.P. and L.C.Z.; formal analysis, U.O., K.A.P. and L.C.Z.; investigation, U.O. and L.C.Z.; resources, F.U. and L.C.Z; data curation, K.A.P.; writing—original draft preparation, U.O., K.A.P., F.U. and L.C.Z.; writing—review and editing, U.O., K.A.P., V.O., C.A., A.G., J.R. and L.C.Z.; visualization, K.A.P., J.R. and L.C.Z.; supervision, L.C.Z; project administration, L.C.Z; funding acquisition, L.C.Z.

## Acknowledgments

The authors thank Octavio Zambada Moreno for his technical assistance in data visualization. This article is dedicated to the memory of Dr. Juan Ignacio Sarmiento Sanchez.

